# Environmental factors drive release of *Perkinsus marinus* from infected oysters

**DOI:** 10.1101/2020.11.03.366781

**Authors:** Sarah A. Gignoux-Wolfsohn, Matilda S. R. Newcomb, Gregory M. Ruiz, Katrina M. Pagenkopp Lohan

**Author notes:** Corresponding Author: Smithsonian Environmental Research Center, 647 Contees Wharf Road, Edgewater, MD 21037.

## Abstract

Since the discovery of *Perkinsus marinus* as the cause of dermo disease in *Crassostrea virginica*, salinity and temperature have been identified as the main environmental drivers of parasite prevalence. However, little is known about how these variables affect the movement of parasites from host to water column. In order to elucidate how environmental factors can influence the abundance of this parasite in the water column, we conducted a series of experiments testing the effects of time of day, temperature, and salinity on release of *P. marinus* cells from infected oysters. We found that *P. marinus* cells were released on a diurnal cycle, with most cells released during the hottest and brightest period of the day (12:00-18:00). Temperature also had a strong and immediate effect on number of cells released, but salinity did not, only influencing the intensity of infection over the course of several months. Taken together, our results demonstrate that 1) the number of parasites in the water column fluctuates according to a diurnal cycle, 2) temperature and salinity act on different timescales to influence parasite abundance, and 3) live infected oysters may substantially contribute to the abundance of transmissive parasites in the water column under particular environmental conditions.

## Introduction

*Perkinsus marinus* is a protistan parasite that infects the eastern oyster (*Crassostrea virginica*) and is notorious for causing dermo disease (Mackin, 1951; Ray, 1954). It was first identified in the 1940s after causing severe mass mortalities of wild oyster populations in the Gulf of Mexico (Ray, 1954). Its current geographic range spans the east coast of the Americas, from northern South America (da Silva *et al.*, 2013), to the Gulf of Maine (Cook *et al.*, 1998), including Central America (Pagenkopp Lohan *et al.*, 2016), and the Gulf of Mexico (Ray, 1954). In the wild, infective *P. marinus* cells are taken into the oyster through filter feeding, with many infections occurring through the mantle from infected pseudofeces (Allam *et al.*, 2013; Ben-Horin *et al.*, 2015). While *P. marinus* epizootics have historically resulted in the collapse of entire oyster reefs (Hewatt & Andrews, 1954), enzootic populations of the parasite are currently common (Calvo *et al.*, 2003). Low-level infections can be sustained for years with only slight effects on growth and reproduction (Ray, 1954; Sunila & LaBanca, 2003), with parasites building up over time in older oysters (Andrews & Hewatt, 1957).

On oyster reefs, *P. marinus* prevalence, intensity, and resulting oyster mortality increases with temperature and salinity (Andrews, 1988; Ray, 1954). Epizootics are therefore seasonally cyclical, with the highest prevalence and intensity in summer and fall months (Andrews & Hewatt, 1957) and lowest in winter months (Andrews, 1965). The severity of outbreaks also varies across years, with particularly intense outbreaks after La Niña events, when both temperature and salinity increase (Soniat *et al.*, 2005). Geographic variation in temperature and salinity also drive *P. marinus* distribution, with warming sea temperatures likely contributing to the northern range expansion of *P. marinus* (Cook *et al.*, 1998; Ford & Chintala, 2006). In the lab, Chu and colleagues found that both higher temperature (Chu & La Peyre, 1993) and salinity (Chu *et al.*, 1993) increased infection prevalence and intensity in oysters challenged with *P. marinus.* In the same experiments, temperature and salinity also altered the presumed cellular defenses of the hosts, such as abundance of hemocytes, suggesting a potential impact on host immune processes. In culture, high temperature and salinity increased development of *P. marinus* hypnospores (Chu & Greene, 1989b) as well as metabolic activity and proliferation of trophozoites (La Peyre *et al.*, 2008), with salinities below 7 ppt causing severe reductions in viability (La Peyre *et al.*, 2006).

In the wild, *P. marinus* is believed to be primarily transmitted between oysters through the water column (Ray, 1954). The parasite is able to survive for weeks outside the host due to its ability to synthesize fatty acids, facilitating both short and long distance (*i.e.,* reef-to-reef) transmission (Chu *et al.*, 2002). Transmission can also occur through ectoparasitic snails feeding on infected oysters (White *et al.*, 1987). Live *P. marinus* cells have been found in the stomachs of multiple species of snails and fish, and on the setae of mud crabs (Andrews *et al.*, 1962), although the relative frequency of transmission by animal vector vs. through the water column is unknown. Even with continuous exposure, detectable infections develop slowly, likely because a high infective dose is required (Bidegain *et al.*, 2016). While timing can vary, naive oysters placed in water where the parasite is endemic or adjacent to heavily infected oysters will generally develop visible infections over the course of weeks (Andrews, 1965; Ray, 1954). For this reason, understanding the factors that contribute to the abundance of infective particles in the water column is crucial to understanding disease dynamics, including rates of infection.

The majority of infective *P. marinus* in the water column is historically believed to come primarily from the death of heavily infected oysters, as the abundance of free-living parasites is highly correlated with oyster mortality (Audemard *et al.*, 2006; Calvo *et al.*, 2003). Large numbers of cells are released from dead and dying oysters, especially when tissues are shredded by scavengers, providing a logical mechanism for these observed patterns (Hoese, 1962). However, death of heavily infected oysters is not the only avenue for the reintroduction of *P. marinus* cells to the water column, and live cells can be released from infected oysters in pseudofeces (Bushek *et al.*, 2002). In the wild, parasite abundance is not only correlated with mortality but also with weighted prevalence in live oysters, suggesting that heavily infected live oysters may also contribute significantly to *P. marinus* abundance in the water (Audemard *et al.*, 2006). In addition, naïve oysters can pick up infections from the water even when oyster mortality is low (Calvo *et al.*, 2003). Because infection and development of disease signs happen over the course of weeks and months, studies of *P. marinus* abundance in the water column have focused on these time scales (Audemard *et al.*, 2006; Calvo *et al.*, 2003). However, on shallow tidal oyster reefs, environmental parameters such as temperature and salinity often fluctuate over much shorter timescales. Little is known about how short-term changes in these parameters affect the release of *P. marinus* cells from live oysters. Here, we conducted two experiments to examine the effects of time of day, salinity, and temperature on the release of *P. marinus* from infected oysters. We found that both time of day and temperature strongly influenced the number of cells released. Further, we found that while generally more cells were released from dead or dying oysters than from live oysters, similar numbers of cells can be released from live oysters at high water temperatures. We surmise that live oysters contribute significantly to the population of *P. marinus* in the water column and should be considered in transmission dynamics.

## Materials and Methods

### Experimental oysters

In order to measure the release of *Perkinsus marinus* by infected oysters, we obtained 200 wild oysters (*Crassostrea virginica*) from the Nanticoke River, Maryland in July 2019. This location was chosen due to previous findings of high prevalence, oysters from this river had a 73% prevalence in 2018, with one of the highest recorded mean intensities in the Maryland portion of the Chesapeake Bay (2.3 on the Mackin scale) (Tarnowski, 2018). Oysters had a mean two-dimensional area of 76.46 (SD 20.82) cm and a mean wet weight (including shell) of 237.21(SD 92.46) g. Prior to the start of the experiments, we sacrificed 55 oysters and measured *P. marinus* infection using Ray’s Fluid Thioglycollate Method (RFTM) as described below. We found a prevalence of 43% and mean intensity of 0.75 (SD 1.16). The remaining oysters were randomly divided into three groups kept in Rhode River water at different salinities: low (5-10 ppt), medium (10-15 ppt), and high (20-25 ppt) manipulated by adding freshwater or aquarium salt (Instant Ocean, Blacksburg VA) as needed (Figure 5A). These levels were chosen to mimic the extremes of the salinity range found in the Maryland portion of the Chesapeake Bay. All oysters were fed Frozen Shellfish Diet for Broodstock (Reed Mariculture, Campbell CA) until the start of Experiment #1. The authors assert that all procedures contributing to this work comply with the ethical standards of the relevant national and institutional guides on the care and use of laboratory animals.

### Experiment #1: Effect of salinity and time of day on P. marinus release over 24 hours

In order to test the effect of salinity on release of *P. marinus* cells over a short period of time, we took oysters (N=36) from the medium salinity holding treatment and placed them in Rhode River water adjusted to three salinity levels, low (5ppt), medium (15ppt), and high (25ppt) as above (Figure 5A). Water was filtered through a 1um mesh filter to ensure that no *P. marinus* cells were introduced via the water. Oysters were acclimated to their experimental salinity levels over the course of two days. At the start of the experiment, (24:00 on August 14, 2019), oysters were placed in individual 1L containers with water and an air stone. To ensure continuous filtration by the oysters, the pre-filtered water was supplemented with a small amount of dilute shellfish diet (Ehrich & Harris, 2015). The mesocosms were placed outside of the wetlab at the Smithsonian Environmental Research Center (SERC) in the shade in order to mimic the change in water temperature over a 24 hour period. To monitor the temperature throughout the experiment, HOBO sensors were randomly placed in three containers and began temperature measurements at 0:00. The water was sampled as described below and replaced with new water every 6 hours, resulting in four sampling points (Figure S1A). All oysters were sacrificed for RFTM at the end of the experiment.

### Experiment #2: Effect of temperature on P. marinus release

In order to test the long-term effects of salinity on *P. marinus* release, the surviving oysters in the high (n=35) and low (n=36) salinity holding treatments were kept for 2 months before the start of Experiment #2 in 300 gallon tanks where water was changed and salinity was adjusted every 3 days. Oysters were labeled with bee tags (Better Bee Greenwich, NY) in order to identify individuals. To measure short term temperature impacts on *Perkinsus* release, each oyster was placed in an individual 1L container with filtered water for a 6 hour period on four separate days (hereafter referred to as trials). Individual containers were placed in a communal water bath to ensure consistent temperatures across replicates. Oysters were kept in communal holding tanks inside at appropriate salinity levels and ambient temperature between trials with timed artificial lights. All trials ran from 9:00 to 15:00 inside the wetlab at SERC under artificial light (Figure S1B). At the conclusion of each 6 hour trial, the oyster was removed and the entire contents of the container was sampled, including water and any particles, feces, and pseudofeces. All oysters experienced a total of four trials: two trials where the water temperature was held at ambient temperatures (25-28 °C) followed by two trials where the water bath was heated (30-32 °C) (Figure 5B). Due to space constraints, oysters were divided into two groups and trials were run on separate days. For heated treatments, temperature was slowly ramped up and down over the course of 24 hours to not shock the oysters. For a given group, trials were always separated by a minimum of 3 days.

### Sampling and analysis

In order to count the number of pathogens released into the water column in both experiments, the 1L container was shaken and 500mL was removed and filtered through a 0.45 μm whatman filter (Cytiva Life Sciences Marlborough, MA) using a glass filter apparatus that was rinsed with bleach in between samples. To measure oyster infection at the end of each experiment, oysters were sacrificed and a piece of anal and mantle tissue was placed in RFTM media. We incubated the filters and tissue in Ray’s Fluid Thioglycollate Media (RFTM) at 27 °C for five to seven days. RFTM media causes viable *Perkinsus sp.* cells to enlarge making them visible under a dissecting microscope (Dungan & Bushek, 2015). We placed tissue or filters on a microscope slide and stained with 2-5 drops of Lugol’s iodine (Sigma Aldrich St. Louis, MO). Staining with Lugol’s solution allows visualization of cells under a dissecting microscope. For tissue samples, we utilized the Mackin Scale, which quantifies infection intensity on a scale from 1 to 5 (Ray, 1954). For filters where total number of cells was less than ~500, we counted all cells. For the twelve filters with a high abundance of cells (>500), we took 8 pictures of random locations on the filter at 4x magnification and counted the number of cells using IMAGEJ (Schindelin *et al.*, 2012). At 4x magnification the filter contains 56 frames, so we multiplied these numbers by 9 to estimate the total number of cells across the entire filter.

### Statistics

We looked at the effect of salinity in Experiments #1 and #2 on number of cells released (log transformed) using ANCOVAs with Mackin score of tissue as a covariate. We conducted a logistic regression with Mackin score as an ordinal response using the poor function in the R package MASS (Ripley *et al.*, 2013) to measure the effect of salinity on intensity of oyster infection for Experiment #2. We used separate GLMMs with a poisson distribution and “replicate” as a random effect in the R package lme4 (Bates *et al.*, 2018) to look at the effect of time of day, temperature, and sunlight on the number of cells released in Experiment #1. To look at the effect of temperature in Experiment #2 for a given oyster, we first averaged the number of cells observed in the two trials (for each oyster, two ambient and two heated). We then subtracted the ambient average from the heated average. In order to transform the data, we added 105 to every value to make all values positive and conducted a boxcox transformation in the r package MASS using a lambda of 0.033. We then conducted a one sample t-test with a null hypothesis of no difference. Finally, to compare the number of cells released by survivors and non-survivors, we first took the maximum of all available values then conducted a glm with a gaussian distribution, with “treatment” and “survival” as factors. We conducted a tukey HSD test using the R package multcomp. All graphs were made using ggplot2 (Wickham, 2009).

#### Data availability

All analyses are available on GitHub https://github.com/sagw/DE2019. Raw will be made available on Figshare.

## Results

### Salinity

In Experiment #1, where oysters were exposed to three salinity levels for three days, salinity had no effect on number of *P. marinus* cells released (ANCOVA DF=6,130, p=0.17). In Experiment #2, where oysters were held at two levels of salinity for four months, salinity also had no effect on number of cells released (ANCOVA DF =2,45 p=0.08). Salinity did, however, have a significant effect on the final intensity of *P. marinus* infections within oyster tissues—while all oysters were infected (100% prevalence), oysters in the high salinity treatment (15-25 ppt) had significantly higher intensity infections at the end of this period (Figure 1, logistic regression p=7.47e-06).

**Figure 1.**
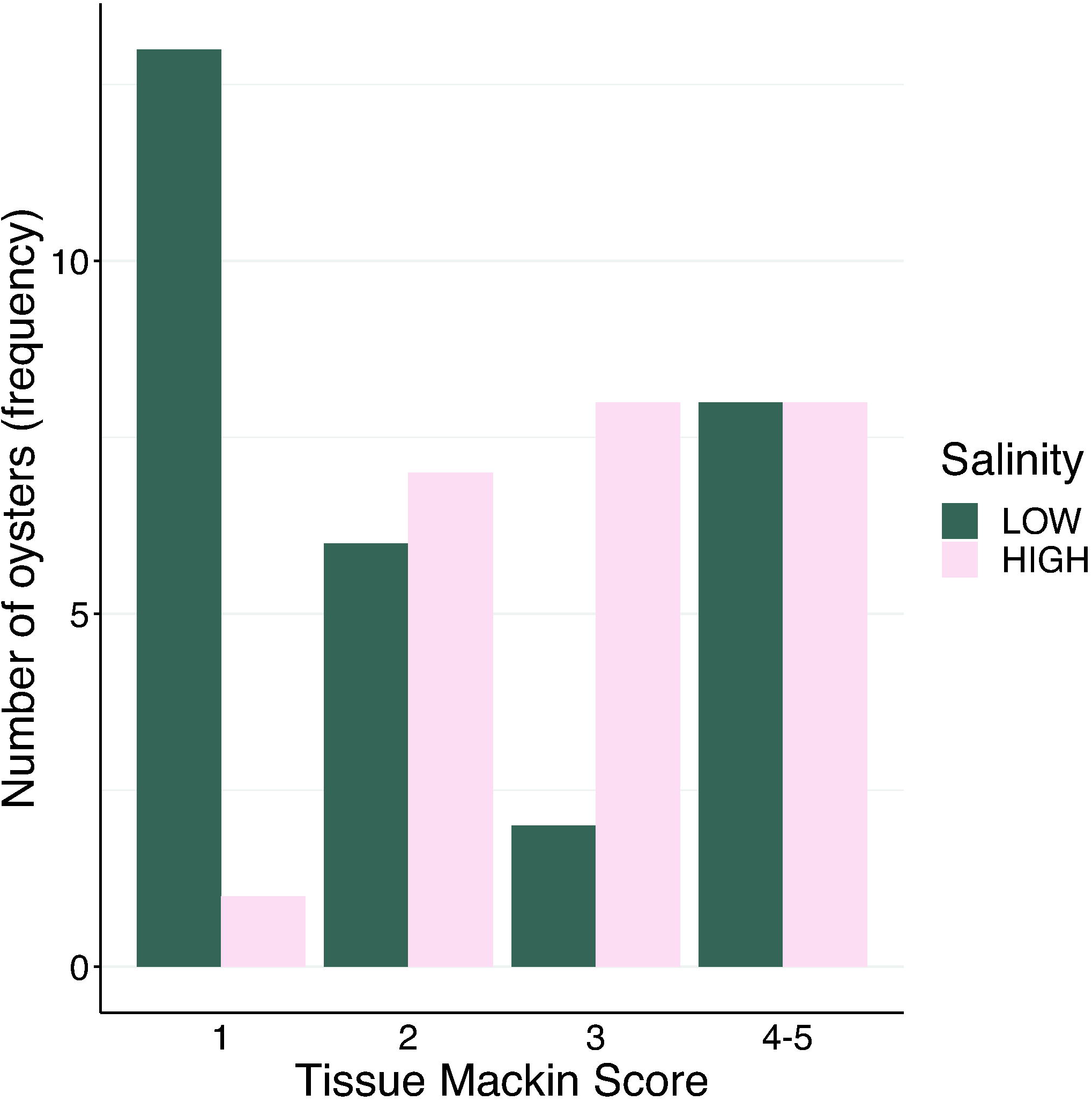
Salinity treatments affected the final intensity of infection of oyster tissues. Histogram showing the number (frequency) of oysters from Experiment #2 with each Mackin score broken up by salinity treatment (across temperature treatments). We combined 4 and 5 due to the few number of 5s observed. Low salinity water was maintained between 5 and 10 ppt and high salinity water was maintained between 20 and 25 ppt.

### Diurnal cycle

Oysters used in Experiment #1, where water was sampled four times over a 24 hour period, had a prevalence of 39% (14/36) and mean intensity of 3.18. *P. marinus* cells were released from nine oysters during at least one of the four sampled time points, with six oysters releasing cells at two timepoints. Cells were only released from infected oysters (*i.e.,* those with a Mackin score greater than 0). Because there was no effect of salinity on number of cells released, salinity was excluded from further analyses. Across all individuals, we observed a diurnal cycle of *P. marinus* release, with the greatest number of cells released between 12:00 and 18:00. For seven out of 9 oysters, the majority of parasite cells were released during this time period (Figure 2). Both temperature (glmm, p<2e-16) and inferred sunlight based on the angle of the sun (glmm, p<2e-16) had significant effects on number of *P. marinus* cells released, with both variables peaking at 18:00 (Figure 2, Figure S2)

**Figure 2.**
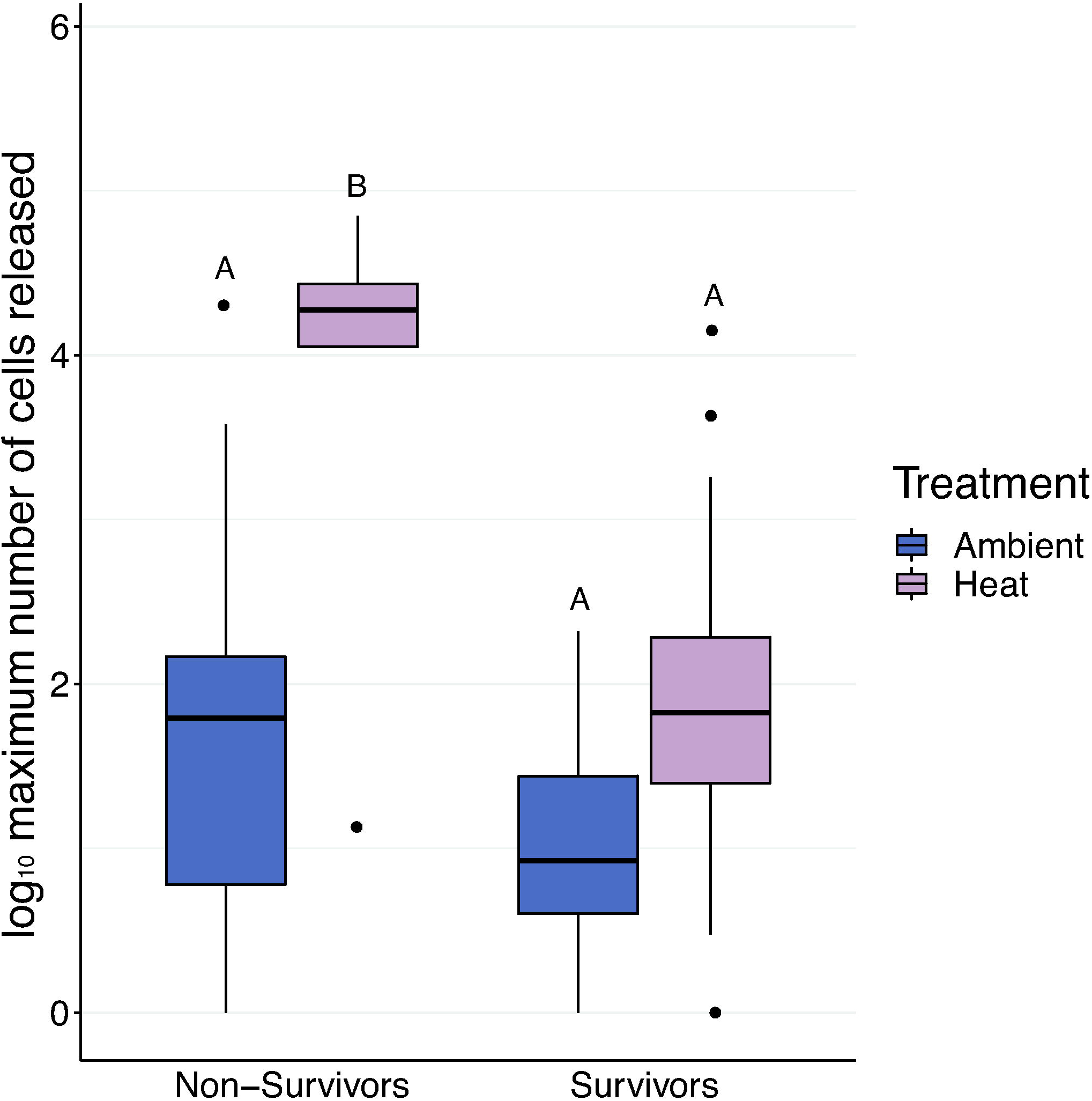
Time of day drove *P. marinus* release. Nine of the oysters used in Experiment #1 released cells. Number of cells found during a timepoint is colored by oyster replicate and points are connected by lines. The time when the sample was taken is shown, meaning that cells were released in the six hour period beforehand. Temperature (°C) as measured by a hobo datalogger is displayed on the right y axis and denoted by triangles.

### Temperature

In Experiment #2, we compared the number of parasite cells released when oysters were in ambient (23–25°C) vs. heated (30–32°C) water temperatures. For 76% (36 out of 47) of the oysters that survived to the end of the experiment, more *P. marinus* cells were released in heated water than in ambient water (Figure 3, one sample t-test, df=46, p=0.036). Neither salinity nor Mackin score of the oyster influenced this change in release.

**Figure 3.**
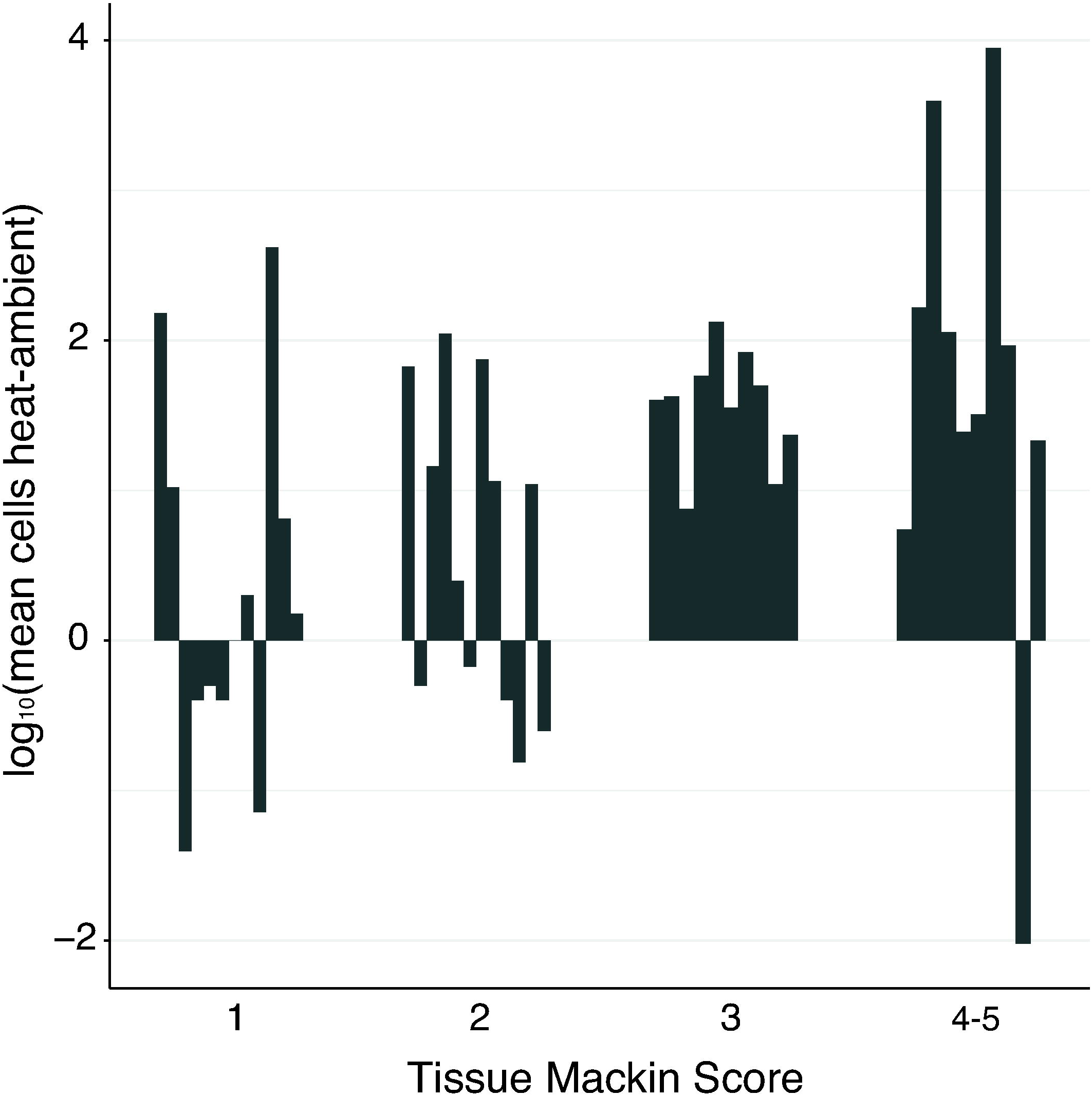
Most oysters released more cells in the heated than ambient treatments. Difference between mean heat and mean ambient cells released for each individual oyster in Experiment #2 shown by bar. Difference was calculated as mean heat cells released-mean ambient cells released for each oyster. Plotted values are the log10 of the absolute value of that difference then adjusted by sign depending on non-adjusted direction of change. (i.e., negative differences remain negative) oysters with no difference are displayed as 0.

### Survival

There were no mortalities in Experiment #1. In Experiment #2, 27% (19 out of 71) oysters died before the conclusion of the experiment and were excluded from the analyses presented above. Salinity did not impact mortality (11 non-survivors were in high salinity and 8 were in low salinity). Fourteen oysters died in the first 20 days of the experiment before being exposed to elevated temperatures and two oysters died after being exposed to one heated trial. We were only able to collect tissues from five of the oysters that died, four of which had a Mackin score ≥ 4. In order to compare cells released from survivors and non-survivors, we calculated the maximum number of cells for each oyster and compared across treatments in order to eliminate issues with number of datapoints (e.g., oysters that died after one trial compared to oysters that survived all four). For 76% percent (39 out of 51) of survivors, the maximum number of cells were released during a heated trial. The highest maximum observed was by a non-survivor in a heated trial (70,173 cells, Figure 4). Overall, non-survivors in heated treatments had significantly higher maxima than all survivors as well as non-survivors in ambient treatments. (Figure 4, Glm, interaction p<0.001, Tukey HSD). However, the highest maximum for survivors (14,247 cells released in a heated treatment) was higher than the maxima of 14 non-survivors.

**Figure 4.**
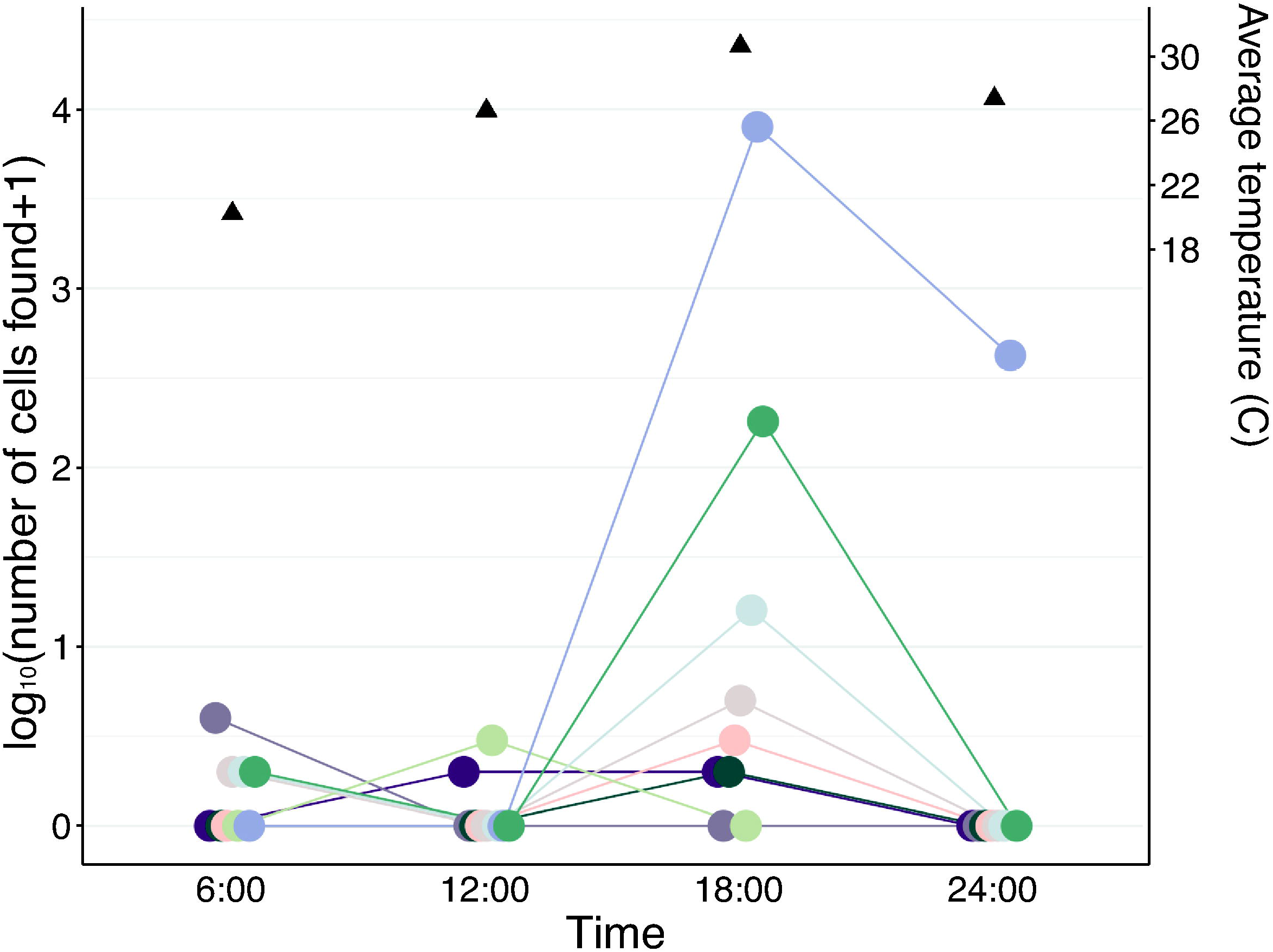
Heated non-survivors released the highest maximum number of cells. Boxplots display mean maxima for a given treatment with outliers are marked by dots. Letters are results of Tukey HSD post-hoc test.

## Discussion

Release of *P. marinus* cells from infected oysters determines the abundance of infectious cells in the water column and likelihood of new infections. Variation in the rate of release can therefore drive overall disease dynamics, particularly within reefs. We build upon previous work showing that viable infective *P. marinus* cells can be released from live infected oysters in feces and pseudofeces (Bushek *et al.*, 2002). By sampling all water surrounding the oyster including (but not limited to) any feces and pseudofeces released during the experiment, we were able to more reliably estimate the number of cells released by a given oyster over time. We examined the influence of environmental factors on rate of release and directly compared the number of *P. marinus* cells released from live to dead and dying oysters. We find that time of day and temperature, but not salinity, have strong effects on the number of cells released in a six hour period (Figures 2 and 3). Because oysters live in estuaries, where temperature and salinity change on daily to yearly cycles, it is important to understand how these factors influence the parasite abundance in the water column on different time scales and the consequences for transmission and disease dynamics in host populations. Our results suggest temperature may be an important driver of parasite abundance in the water column at short time scales, in addition to those previously reported for seasonal and annual scales. More broadly, our data demonstrate that large numbers of parasite cells were released from both live and dead oysters, suggesting that live oysters should not be discounted as significant contributors to *P. marinus* abundance in the water column (Figure 4).

Salinity has consistently been identified as an important driver of *P. marinus* distribution and dynamics (Mackin, 1951; Ray, 1954; Soniat *et al.*, 2005). In the field, naive oysters placed in regions with high salinity developed higher intensity infections than those placed in regions with low salinity (Paynter & Burreson, 1991), either due to higher abundances of *P. marinus* in the water column leading to increased infection pressure, higher pathogenicity, and/or a higher reproduction rate within the oyster. When oysters are inoculated with a standard number of *P. marinus* cells and placed in a closed system, higher salinity increases the intensity of infections over the course of several weeks, suggesting that the main effect of salinity is on *P. marinus* reproduction within the oyster host (Chu *et al.*, 1993; Fisher *et al.*, 1992; Ragone Calvo & Burreson, 1993). Our result that oysters in the high salinity treatment had higher intensity infections after four months corroborates this finding—while we did not use a fully closed system, water in the holding tank was changed infrequently (biweekly) and we therefore assume that the vast majority (if not all) of new infections were acquired from neighboring oysters rather than the incoming water. Together, these findings suggest that the main effects of salinity on *P. marinus-C. virginica* interactions are on the increased abundance of the parasite inside the oyster rather than on increased infectivity or behavior in the water column. Whether this effect is due to a reduction of oyster defense/immunity or an increase in parasite reproduction rate due to higher parasite fitness should be further investigated. Both of our experiments demonstrated that when infection intensity is accounted for, salinity does not impact the rate at which an *P. marinus* cells are released over the course of several hours. Therefore, increased abundance of *P. marinus* in high salinity water is likely due to an increase in the intensity of infections resulting in higher numbers of *P. marinus* cells released into the water column. While *in vitro* low salinity (7ppt) has been shown to reduce viability of *P. marinus* (La Peyre *et al.*, 2006), we did not see an impact of the low salinity treatment on the number of *P. marinus* cells released in either experiment. This could be because our low salinity treatment was not low enough to reduce viability, or because those cells released in the low salinity treatment would not have been viable if they remained at that salinity for a longer period of time rather than being put into RFTM media after 6 hours.

Our observed diurnal cycle of *P. marinus* release was correlated with both water temperature and solar exposure, factors that could influence both host and parasite processes. Oysters have been shown to display both daily and seasonal cycles of feeding activity, but this is the first study to look for daily cycles of *P. marinus* release into the water column. Work on *Crassostrea gigas* has confirmed the presence of an endogenous circadian rhythm that can be influenced by season (Mat *et al.*, 2012), light, and tidal cycle (Tran *et al.*, 2011). This circadian rhythm could be driving oyster feeding and defecation and therefore the release of *P. marinus* cells observed here. Temperature is also a strong driver of oyster behavior and interacts with these other influences on circadian rhythm. Early experiments by Galtsoff on *C. virginica* from the Chesapeake bay found the rate of water intake is dependent on temperature, with oysters remaining closed below temperatures of 5°C (Galtsoff, 1928). These results have been replicated by Comeau 2012 who found that *C. virginica* from the Gulf of St. Lawrence increase the number of valve openings (and therefore water filtration) as water temperatures warm in the spring. Interestingly, however, once oysters entered an “active” period in the spring, temperature no longer exerted an influence on opening, with oysters staying open for periods of time ranging from 1 minute to 13.8 days. During the active period, oysters did exhibit a diurnal cycle independent of temperature, with both length of time and degree of valve openness peaking at 18:00 hours (Comeau *et al.*, 2012). Time of day alone could therefore explain our finding that maximum release of *Perkinsus* cells occurred between 12:00 and 18:00 hours irrespective of temperature. However, our oysters and *P. marinus* also experienced a much greater fluctuation in temperatures over a 24 hour period than the oysters studied by Comeau *et al.,* and these more drastic changes in temperature likely influenced *P. marinus* release in addition to time of day.

Experiment #2 allowed us to test the effects of temperature on *P. marinus* release regardless of time of day, demonstrating that temperature alone increases the rate of *P. marinus* release. Temperature is known to be an important driver of *P. marinus* reproduction and infectivity both *in vitro* (Chu & Greene, 1989a; Chu & Volety, 1997) and *in vivo* (Calvo *et al.*, 2003) (Fisher *et al.*, 1992), and could therefore alter pathogen behavior or reproduction leading to increases in release. By counting the number of *P. marinus* cells released from a given oyster in both ambient and heated environments, we controlled for the significant individual variation observed in Experiment #1 and the strong influence of infection intensity on number of cells released. This experimental design allowed us to get reliable estimates of the increase in number of cells released due to increased temperature. The water temperature fluctuations in Experiment #1 were consistent with temperatures experienced by intertidal oysters in shallow embayments of the Chesapeake Bay during summer months (Maryland Department of Natural Resources). In the wild, both mean temperature and temperature variability will be highly dependent on location within the estuarine environment and relation to tidal height (i.e., subtidal vs. intertidal and amount of time spent out of water). Previous work has found that intertidal oysters have higher prevalence and intensity of *Perkinsus* infections than subtidal oysters (Malek & Byers, 2017), with a positive correlation between intensity and air temperature (Malek & Byers, 2018), indicating that the higher temperatures experienced when out of the water contribute to intensity. Some of the spatial heterogeneity of *Perkinsus* abundance in estuaries is likely driven by a combination of spatial and temporal variability in temperature (Malek & Breitburg, 2016). In order to accurately capture the maximum *P. marinus* abundance in the water column at a given location, water sampling should occur in the afternoon close to the hottest part of the day.

While temperature and salinity are often cited as the two main environmental factors that influence *P. marinus* abundance and infectivity, our results show that they operate on different components of the *P. marinus-C. virginica* interaction and at different timescales. Further work is needed to understand when and how temperature and salinity influence the host and parasite individually resulting in the patterns observed here and by others. Controlled and fully crossed experiments are needed to tease out the interactive effects between light, time of day, and temperature, which likely influence both the host and parasite in ways not fully captured by our experiments. In addition, future experiments should 1) keep temperature more tightly controlled 2) examine a wider range of temperatures and 3) alter the length of the experiments to test the effect of duration of temperature changes on release of parasite cells.

Seven pathogenic species of *Perkinsus* have been identified to date, infecting a wide variety of hosts (Villalba *et al.*, 2004). While *P. marinus* is the dominant species in Chesapeake Bay populations of *C. virginica, P. chesapeaki* has also been found in these oysters, albeit at much lower prevalence and intensity (Reece *et al.*, 2008). We assumed that the *Perkinsus* cells detected here using RFTM were *P. marinus,* based on previous work showing this is the dominant parasite in oysters of this region (Gignoux-Wolfsohn unpublished data, (Reece *et al.*, 2008). Future work should use molecular methods to both identify the species released from experimental oysters and test if *P. chesapeaki* and other species exhibit similar rates of release to *P. marinus.*

Previous research demonstrated that infective *P. marinus* cells are released through feces and pseudofeces of live oysters (Bushek *et al.*, 2002). However, this release was assumed to be negligible when compared to release by the scavenging and disintegration of infected dead oyster tissue (Audemard *et al.*, 2006; Calvo *et al.*, 2003). We found that while more cells are generally released from dead and dying oysters than live oysters, live oysters are capable of releasing tens of thousands of cells over the course of 6 hours and should therefore be considered a major contributor to the number of parasites in the water column. Given our finding that live oysters release more cells at higher temperatures, the relative contributions that live and dead oysters make to parasites in the water column may be heavily temperature dependent. In this experiment, the cause of death was not always obvious as many oysters died before tissue could be collected for RFTM. However, four out of the five non-survivors that were assayed had heavy infections suggesting that the parasite played a role in mortality. While release from dead and dying oysters may result in a spike of *P. marinus* cells in the water, live oysters have the potential to contribute to a steadier release of cells for a longer period of time. Seven of the survivors released cells at all four timepoints assayed. Furthermore, infective cells were consistently released from live oysters with low intensity. Once a dead oyster has disintegrated or been eaten, it no longer acts as a source for the water-borne parasite population, whereas live oysters may contribute to this population to varying degrees for an entire season. Over the course of several years, live oysters harboring low-level infections could result in the release of equivalent or larger amounts of *P. marinus* cells than oysters that quickly develop a high-intensity infection and die in one season. Live infected oysters should therefore be considered as a significant contributor to enzootic *P. marinus* populations in the water column.

In summary, we measured the effects of environmental factors and time of day on release of *P. marinus* cells from *C. virginica.* We found that temperature and time of day had strong effects on the number of parasites released, but that salinity only influenced the abundance of the parasite within its host. Furthermore, we found that live oysters can release numbers of *P. marinus* cells that are comparable to the amount released from dead and dying oysters. These factors should all be taken into consideration when estimating the population of free-living *P. marinus* in the water column and variations in this population over both hourly and seasonal timescales.

## Figure Legends

Figure S1. Experimental set up and experiment timeline. A) Experiment #1. Each oyster represents a single replicate inside a single tupperware. The oyster remained inside the tupperware for the duration of the experiment, as water was removed and replaced. B) Experiment #2. Each oyster experienced a total of 4 trials, but only half of the oysters were assayed on a given day due to space restraints. The two days under the trial number indicate when that trial was conducted (half the oysters assayed on the first day and half on the second). Oysters were removed from tupperware and placed in communal holding tanks in between trials.

Figure S2 Time of day drove *P. marinus* release. Nine of the oysters used in Experiment #1 released cells. Number of cells found during a timepoint is colored by oyster replicate and points are connected by lines. The time when the sample was taken is shown, meaning that cells were released in the six hour period beforehand. Sunlight (kilowatt hours/m2) as inferred from the angle of the sun is displayed on the right y axis and denoted by exes.

## Acknowledgements

We would like to thank Kristina Borst and Theresa Vaillancourt for assistance with Experiment #2, George Smith for experiment construction advice, Shannon Hood for help in procuring infected oysters, and Erik Holum for help with automating image analysis.

