## Supplementary figures and images for "Environmental factors drive release of *Perkinsus marinus* from infected oysters"

### Supplemental Figure 1

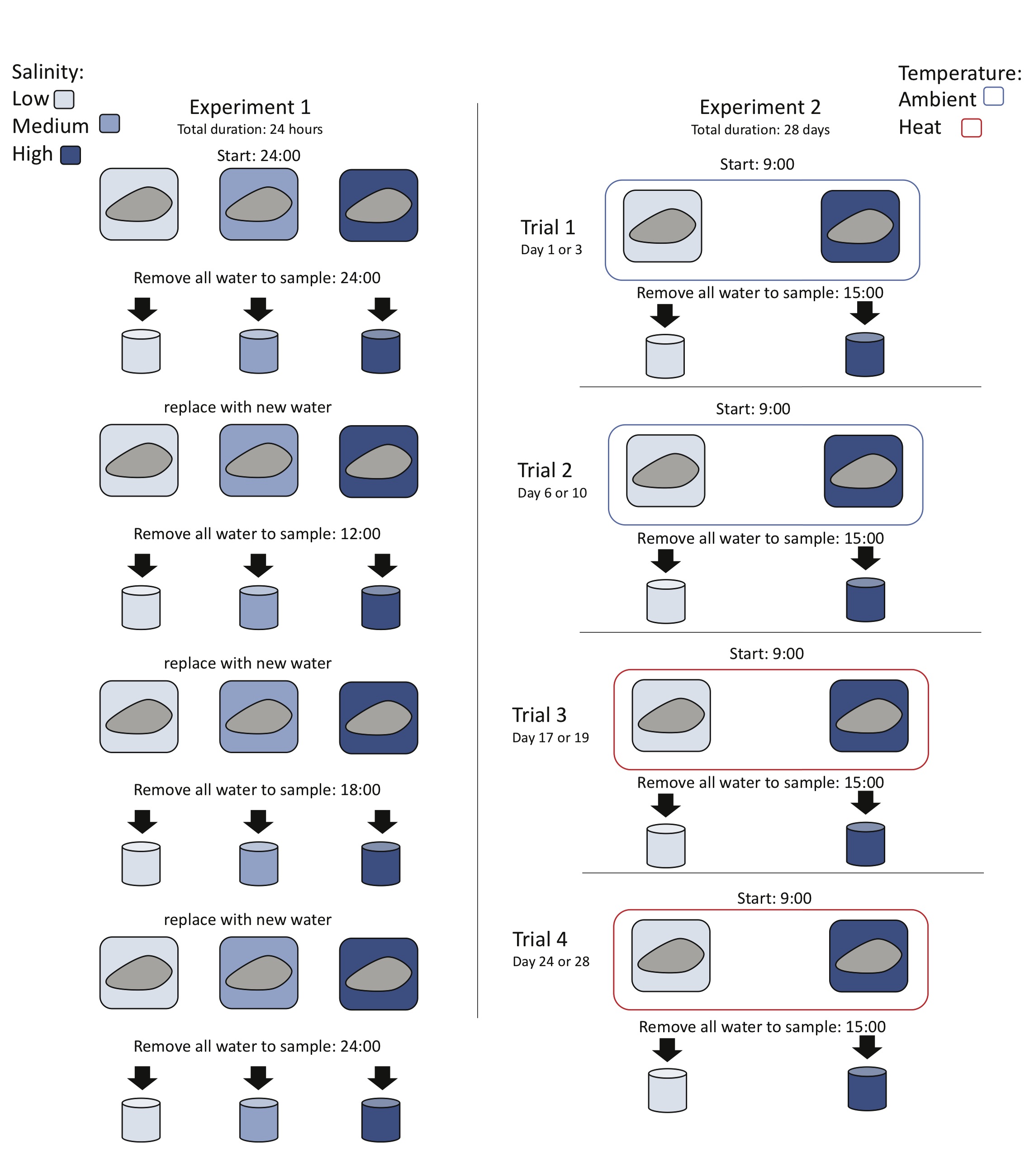

### Supplemental Figure 2

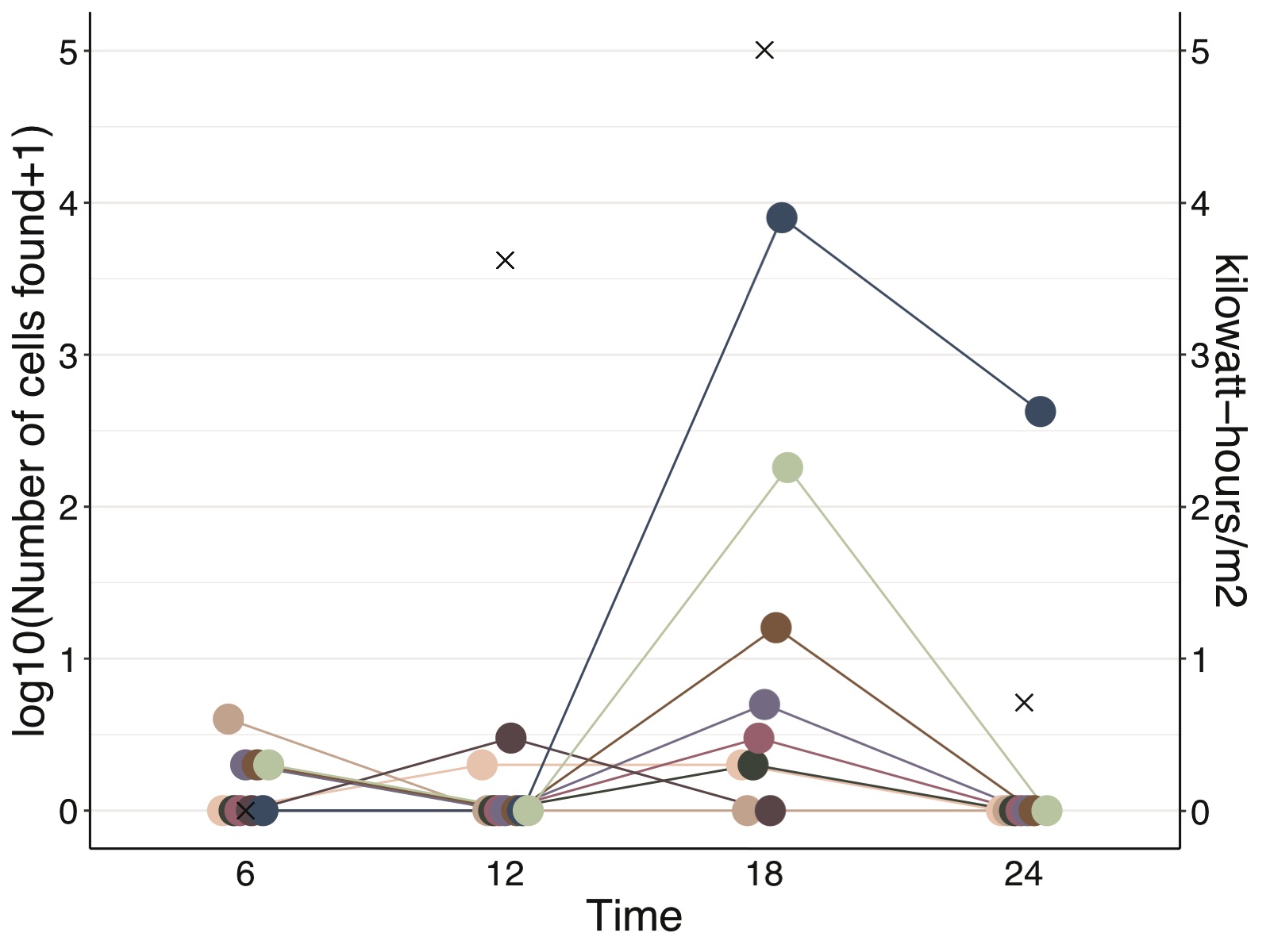
